# Antagonistic effects of actin-specific toxins on *Salmonella* Typhimurium invasion into mammalian cells

**DOI:** 10.1101/2024.07.01.601609

**Authors:** David B. Heisler, Elena Kudryashova, Regan Hitt, Blake Williams, Michelle Dziejman, John Gunn, Dmitri S. Kudryashov

## Abstract

Competition between bacterial species is a major factor shaping microbial communities. In this work, we explored the hypothesis that competition between bacterial pathogens can be mediated through antagonistic effects of bacterial effector proteins on host systems, particularly the actin cytoskeleton. Using *Salmonella* Typhimurium invasion into cells as a model, we demonstrate that invasion is inhibited if the host actin cytoskeleton is disturbed by any of the four tested actin-specific toxins: *Vibrio cholerae* MARTX actin crosslinking and Rho GTPase inactivation domains (ACD and RID, respectively), TccC3 from *Photorhabdus luminescens*, and *Salmonella’s* own SpvB. We noticed that ACD, being an effective inhibitor of tandem G-actin binding assembly factors, is likely to inhibit the activity of another *Vibrio* effector, VopF. In reconstituted actin polymerization assays confirmed by live-cell microscopy, we confirmed that ACD potently halted the actin nucleation and pointed-end elongation activities of VopF, revealing competition between these two *V. cholerae* effectors. Together, the results suggest bacterial effectors from different species that target the same host machinery or proteins may represent an effective but largely overlooked mechanism of indirect bacterial competition in host-associated microbial communities. Whether the proposed inhibition mechanism involves the actin cytoskeleton or other host cell compartments, such inhibition deserves investigation and may contribute to a documented scarcity of human enteric co-infections by different pathogenic bacteria.

## Introduction

Cross-kingdom polymicrobial infections of the digestive system are common and have been seen with nearly every global viral pandemic^1–3^. In contrast, intestinal polybacterial infections are far less common, possibly reflecting a more intense competition between similar organisms occupying the same niche^3–5^. Indeed, complex bacterial communities, such as those that populate the human digestive tract, encompass thousands of species that coexist in highly intricate competitive/cooperative relationships^6^. Of the various competition types, the most recognized are *i)* exploitative competition, when the species compete for essential resources, and *ii)* interference competition, when bacteria directly harm their rivals via effector molecules^6^. The latter is achieved by using secretion systems type I, IV, V, VI, and VII (T1SS, T4-7SS)^7^, which can deliver antibacterial toxins/effector domains that are either secreted to the medium or directly injected into the cytoplasm of the competitor bacteria.

However, it is likely that microbial competition within host is not limited to the above well described mechanisms. Because pathogenic bacteria utilize host cells and extracellular components as a source of nutrients and a niche for growth, we hypothesize that altering the host’s cell pathways and compartments may mediate another, poorly understood and insufficiently explored mechanism of indirect competition between pathogenic bacteria. Specifically, we noticed that while numerous bacterial pathogens target the actin cytoskeleton, they do so for various reasons and in various, sometimes mutually disruptive, ways^8^. By acting directly on actin, actin-binding proteins (ABPs), or related signaling cascades, some pathogens promote actin dynamics to orchestrate membrane remodeling and invade the host cell (*e.g., Salmonella*^9^*, Legionella*^10^). Other pathogens utilize the motoric function of actin polymerization for locomotion inside the host cell (*e.g.*, *Listeria*^11,12^, *Shigella*^12^, *Rickettsia*^11,13^, *Burkholderia*^14^). In contrast, many other effector proteins disrupt actin filaments to compromise the integrity of epithelial barriers, facilitate colonization, and disrupt the phagocytic activity of immune cells (*e.g., Vibrio*^15–17^, *Mycoplasm*^18^, enterohemorrhagic *Escherichia coli* (EHEC)^19,20^). Therefore, we hypothesized that the effects of some pathogens may conflict with the mechanisms employed by other bacterial pathogens, resulting in indirect competition for common host targets such as actin cytoskeleton.

To test the hypothesis that pathogen competition can proceed through manipulating the host systems and, specifically, the actin cytoskeleton, we employed an *in vitro* model of *Salmonella* Typhimurium invasion into immune and non-immune cells pre-treated with various actin-targeting bacterial effectors that impose distinct molecular mechanisms for altering the actin cytoskeleton. Rho GTPase inactivation domain (RID) and actin crosslinking domain (ACD) of the *V. cholerae* multifunctional autoprocessing repeats-in-toxin (MARTX) toxin are liberated inside the host cell by an autoprocessing cysteine protease domain (CPD) and function as separate effectors conferring their individual cytotoxic activities^21^. RID inhibits Cdc42 and other Rho family GTPases by N^ε^-fatty acylation^22^ (*i.e.,* counteracts the activity of *Salmonella* effectors SopB, SopE/SopE2), thereby, favoring actin depolymerization. While employing a very different mechanism, ACD also inhibits global actin dynamics by covalently crosslinking monomeric actin into oligomers^23–25^, which potently inhibit all key actin assembly factors: formins, Ena/VASP, Arp2/3 complex nucleation promoting factors (NPFs), and spire^15,16^. In contrast to F-actin destabilization by RID and ACD, TccC3 domain of a giant ABC toxin from *Photorhabdus luminescens* ADP-ribosylates theonine-143 on actin, which stabilizes F-actin^26,27^, but weakens filaments interaction with the CH-family of actin bundling and network-stabilizing proteins (*e.g.,* plastins and α-actinins)^27^. This, in turn, weakens the integration and connection of the cytoskeleton elements with the cell membrane.

Interestingly, effectors with conflicting activities can even be produced by the same organism. Thus, *S.* Typhimurium both hijacks actin dynamics to invade host cells (by producing SipA, SopB, SopE and SopE2 effector proteins^28^) and destabilizes the actin cytoskeleton (by secreting another effector protein, SpvB, into already infected cells^29,30^). Therefore, we also evaluated the effects on invasion of *Salmonella’s* own effector SpvB. SpvB mono-ADP-ribosylates actin at arginine-177, prohibiting its polymerization^31^, and thus, possibly conflicting with actin-stabilizing effects of SipA and actin dynamics potentiation by SopB and SopE effectors.

Finally, we anticipated and explored another potential interference between *V. cholerae* effectors, VopF and ACD. Such interference can be predicted based on the presence of three tandem G-actin binding WH2 domains in VopF, which is a consensus target for ACD-produced actin oligomers^25^. We tested and confirmed the functional competition between VopF and ACD in reconstituted actin polymerization assays and in live eukaryotic cells.

## Results

### Invasion of Salmonella can be abolished by actin-specific effectors

To test the hypothesis that bacterial competition can be mediated via mutually interfering influences of effectors on the host, we created a model system utilizing *S.* Typhimurium invasion into murine macrophage J774A.1 and epithelial-origin HeLa cells that were pre-treated with various bacterial effectors (Fig. 1). To evaluate the effects of actin cytoskeleton disrupting toxins on the *Salmonella* invasion efficiency, we used constructs containing the amino acid sequences of ACD, RID, TccC3, and SpvB effectors fused at the C-terminus of the anthrax toxin lethal factor N-terminal fragment (LF_N_), which, in cooperation with the protective antigen (PA), allows cytoplasmic delivery of heterologous proteins via the anthrax toxin machinery^32^. These effectors were selected to evaluate the generality of the anticipated interferences as they represent different ways of altering the actin cytoskeleton.

**Figure 1.**
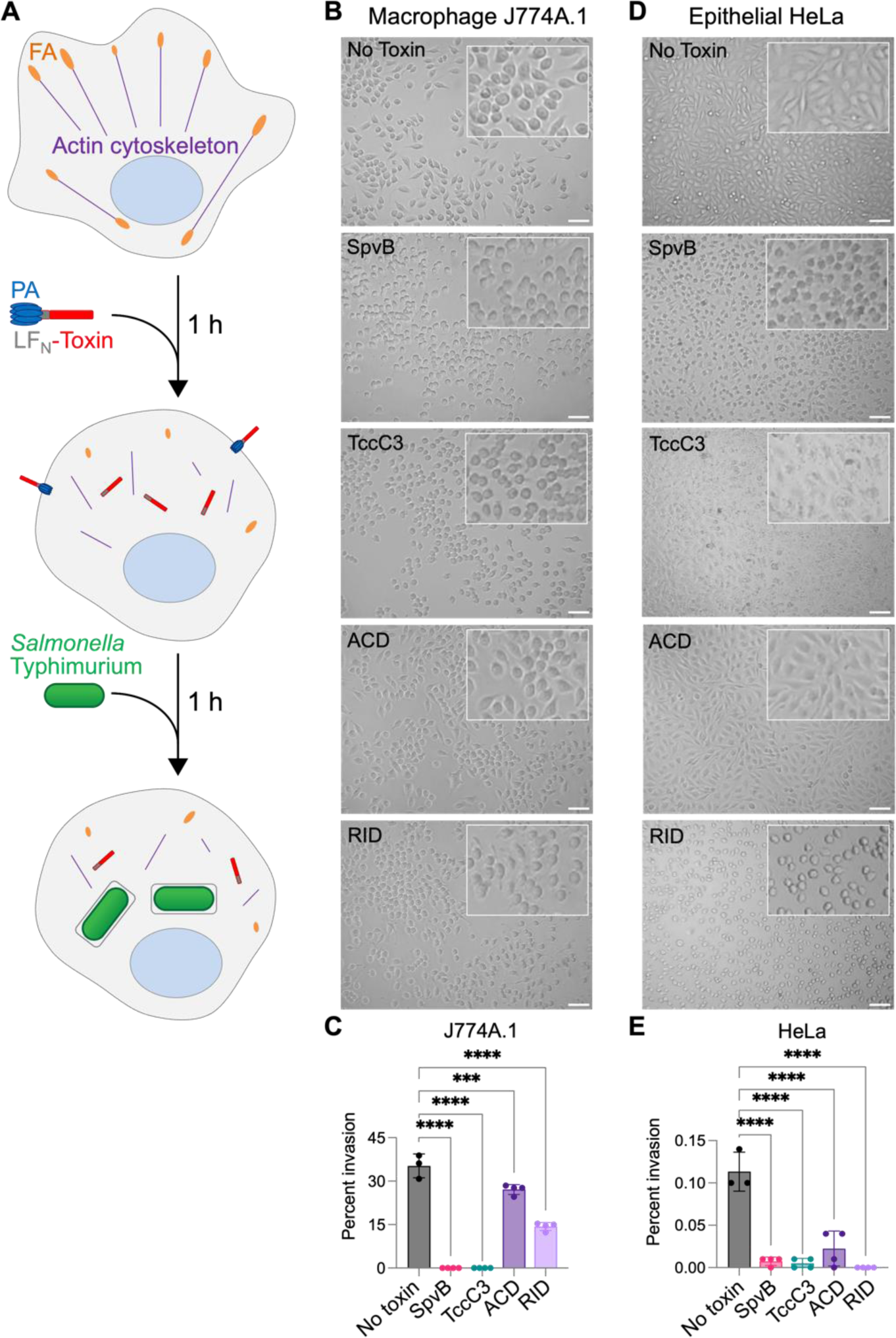
Bacterial actin-specific effectors interfere with *Salmonella* invasion. (A) Schematic diagram shows experimental set up for *Salmonella* invation into J774A.1 and HeLa cells. Cells were pre-treated for 1 h with a mixture of 2.5 nM PA delivery component and 1 nM of a corresponding LF_N_-effector toxin before the addition of *S.* Typhimurium. (B,D) Micrographs of toxin-pretreated J774A.1 (B) and HeLa (D) cells. Scale bars are 50 µm (C,E). *S.* Typhimurium invasion was calculated as described in the Methods section. Data represent the means of 3-4 repetitions ± SD. Ordinary one-way ANOVA with Dunnett’s multiple comparisons test was performed to determine statistical significance; *** = 0.0002, **** < 0.0001.

Macrophage cells are recognized as targets of *Salmonella* that contribute to bacterial spread to distal organs in systemic salmonellosis^33^. To determine if *Salmonella* virulence-plasmid effector SpvB, which mono-ADP-ribosylates actin at Arg177 preventing its polymerization^31^, can affect *Salmonella* invasion, J774A.1 macrophage cells were pre-treated with a mixture of LF_N_SpvB and PA (Fig. 1A,B). As the first morphological changes due to the effector protein entering the cell and disrupting the cytoskeleton by this delivery pathway appear in ∼45 minutes^16,27^, cells were treated for 1 hour and then infected with *Salmonella* (Fig. 1A). After incubation for another 1 hour, extracellular bacteria were killed by gentamicin addition, followed by cell lysis and bacterial numeration^34^. Under these conditions, SpvB reduced the efficiency of *Salmonella* invasion from 35.26% to 0.01%, *i.e.,* rendering it nearly eliminated (Fig. 1C).

To determine whether this phenotype was specific to disrupting the actin cytoskeleton by SpvB^29,30^, we assessed *Salmonella* invasion into macrophage cells treated by TccC3 from *P. luminescens*, which ADP-ribosylates actin at Thr148^26^. This modification stabilizes actin filaments but disrupts their integration into larger assemblies and their interaction with other cellular components^27^. Similar to the effects of SpvB, ADP-ribosylation of actin by TccC3 drastically reduced *Salmonella* invasion from ∼35% to 0.02% (Fig. 1C). To further test the hypothesis that the invasion can be prevented by various types of effector-mediated disruption of actin dynamics, we tested the effects of ACD and RID effector domains of *V. cholerae* MARTX toxin on *Salmonella* invasion. Both effectors inhibit actin dynamics albeit by different mechanisms. While their effects on *Salmonella* invasion were notably weaker compared to those of SpvB and TccC3, both ACD and RID significantly (from 35% to 27% and 14%, respectively) reduced the invasion of J774A.1 macrophage cells by *Salmonella* (Fig. 1C).

We next asked if the disruption of actin dynamics can prevent infection in non-phagocytic epithelial-derived HeLa cells. Similarly to macrophage cells, HeLa cells were pretreated with each of the toxins for 1 h prior to challenging cells with *Salmonella* (Fig. 1A,D). All four toxins potently (from 0.11% to 0.02% or less) prevented *Salmonella* from invading HeLa cells, suggesting that any disruption of the actin cytoskeleton has profound effects on *Salmonella* invasion (Fig. 1E). The more prominent inhibition of *Salmonella* invasion into epithelial cells (Fig. 1E) versus macrophage cells (Fig. 1C) is likely due to the different mechanisms used to invade these cells (*e.g.,* SPI1-effector stimulated changes in actin dynamics in HeLa cells versus active phagocytosis of bacteria by macrophages). Together, these data suggest that bacterial pathogens may disrupt actin dynamics to prevent the host cell invasion by other pathogens and thus obtain a competitive advantage in colonizing the host organism. Alternatively, such invasion inhibition may prevent reinfection of already infected cells by the same bacterial species, as shown here for SpvB or reported previously for competition between *Salmonella* SopE and SptP effectors^35^.

### Functional competition between *Vibrio* VopF and ACD

The production of effectors with countering activities on the actin cytoskeleton is not restricted to *Salmonella*. In addition to the ACD and RID effectors described above that are delivered to the host cells as a part of the T1SS MARTX toxin cassette^36^, some *V. cholerae* strains produce several other actin-targeting effectors. For example, in T3SS-positive strains, the translocated effector VopF is reported to promote intestinal colonization by nucleating actin filaments and disorganizing the actin cytoskeleton^37–39^. Recently we discovered that, in addition to actin nucleation, VopF and its structural homolog (71% sequence homology) from *V. parahaemolyticus* VopL^39,40^ disrupt actin cytoskeleton polarity by promoting unconventional pointed-end processive actin elongation^41^. Both the nucleation and elongation activities of VopF/L require tandem WH2 domains, which are canonical targets of the covalent actin oligomers produced by ACD^15^.

To assess whether ACD inhibits VopF/L activities, we conducted actin polymerization assays in bulk and at the single-filament level (Fig. 2). Bulk pyrene-actin polymerization assay revealed that the enhancement of actin dynamics by VopF and VopL was potently inhibited by ACD-crosslinked actin oligomers, which had an inhibitory concentration (IC_50_) of 3.8 nM and 4.3 nM for VopF (Fig. 2A,B) and VopL (Fig. 2C,D), respectively. These values are consistent with the inhibition measured for mammalian actin assembly factors^15,16^.

**Figure 2.**
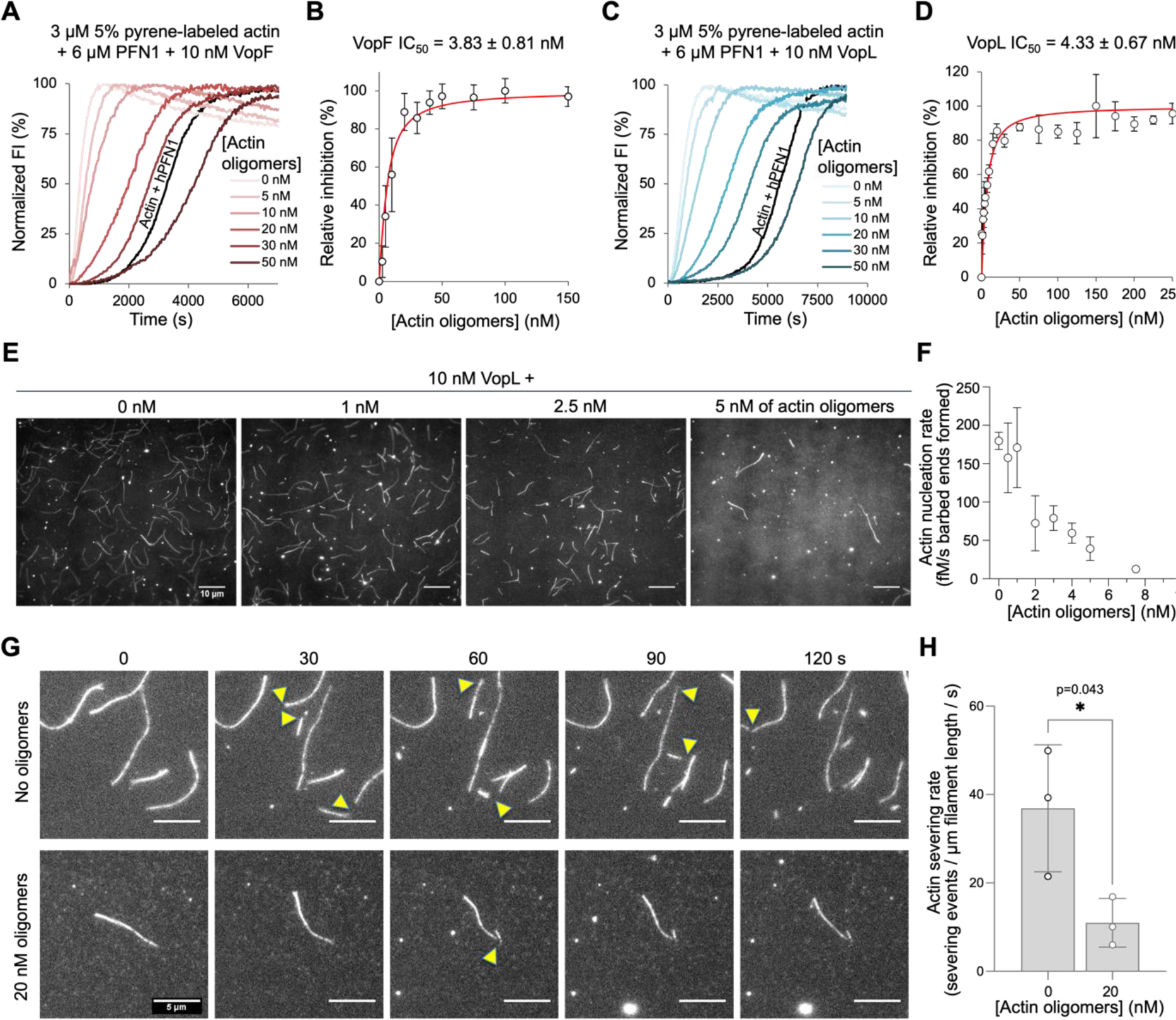
ACD inhibits VopF and VopL actin nucleation and severing activities in the reconstituted assays. (A-D) Effects of ACD-crosslinked actin oligomers on actin polymerization in the presence of PFN1 and VopF (A,B) or VopL (C,D) was monitored in bulk pyrenyl-actin assays (A,C). Normalized FI is pyrenyl-actin fluorescence intensity expressed in percent of maximum. Black traces in A and C is actin/PFN1 polymerization in the absence of VopF/L. Relative inhibition of nucleation by actin oligomers was assessed by the slope at the half maximal fluorescence for VopF (C) or VopL (D) and plotted as means of three experiments ± SD. (E-H) Effects of ACD-crosslinked actin oligomers on actin nucleation and severing by VopL were visualized in TIRF microscopy experiments. (E) Representative images of actin filaments labeled with Oregon Green formed after 10 minutes in the presence of 10 nM VopL and increasing concentrations of oligomers. Scale bars are 10 μm. (F) The total number of filaments in three separate experiments were manually counted and ploted as means ± SE. (G) Representative images of preformed actin filaments severed by VopL in the presence of increasing concentrations of oligomers; time is shown in seconds. (H) Severing rate was calculated from three separate experiments; data is presented as means ± SD, dots are individual data points. Student’s t-test was used to determine statistical significance; **\*** p=0.043.

VopF and VopL potentiate actin dynamics by *i)* nucleating actin filaments^40,42^ and *ii)* promoting the processive filament growth at their pointed end^41^. To assess whether the ACD-produced actin oligomers inhibit nucleation by VopF/L, we counted the number of filaments formed in the presence and absence of ACD-produced actin oligomers using *in vitro* total internal reflection fluorescence (TIRF) microscopy. To minimize the number of spontaneously nucleated actin filaments, profilin (PFN1), a protein that inhibits nucleation of actin filaments, was included in these experiments. In agreement with the bulk polymerization assays, the number of VopL-nucleated actin filaments was significantly reduced with as little as 5 nM of actin oligomers (Fig. 2E, F). The VopF/L severing activity, due to its binding to the filament sides with its WH2 domains and weakening contacts between actin subunits^42^, was similarly inhibited (Fig. 2G,H).

Because the pyrene-actin and standard *in vitro* TIRF assays are reliable reporters of the nucleation and severing activities but not the elongation activity of VopF/L^41^, and to confirm the predicted toxins’ interference in a cellular context, we evaluated the actin-based motility of EGFP-VopF constructs in *Xenopus* fibroblast XTC cells treated with PA/LF_N_-ACD using single-molecule speckle microscopy (SiMS), which reliably reports on the VopF/L processive pointed-end actin elongation activity^41^ (Fig. 3; Movies 1, 2). While the control cells treated with catalytically inactive ACD^43^ showed no significant change in the VopF-controlled filament elongation within cells over the duration of 90 minutes (Fig. 3A,C; Movie 1), the delivery of active ACD reduced the elongation rate by a factor of two within 30 minutes, and resulted in a near-complete halt of pointed-end elongation within 1.5 h (Fig. 3B,D; Movie 2). These effects were notable before changes in cell morphology were detected. Together, these data suggest that the ACD-crosslinked actin oligomers are potent inhibitors of VopF/L, and in strains encoding both proteins, their simultaneous activity could lead to a conflict between ACD and VopF for executing their virulence effects.

**Figure 3.**
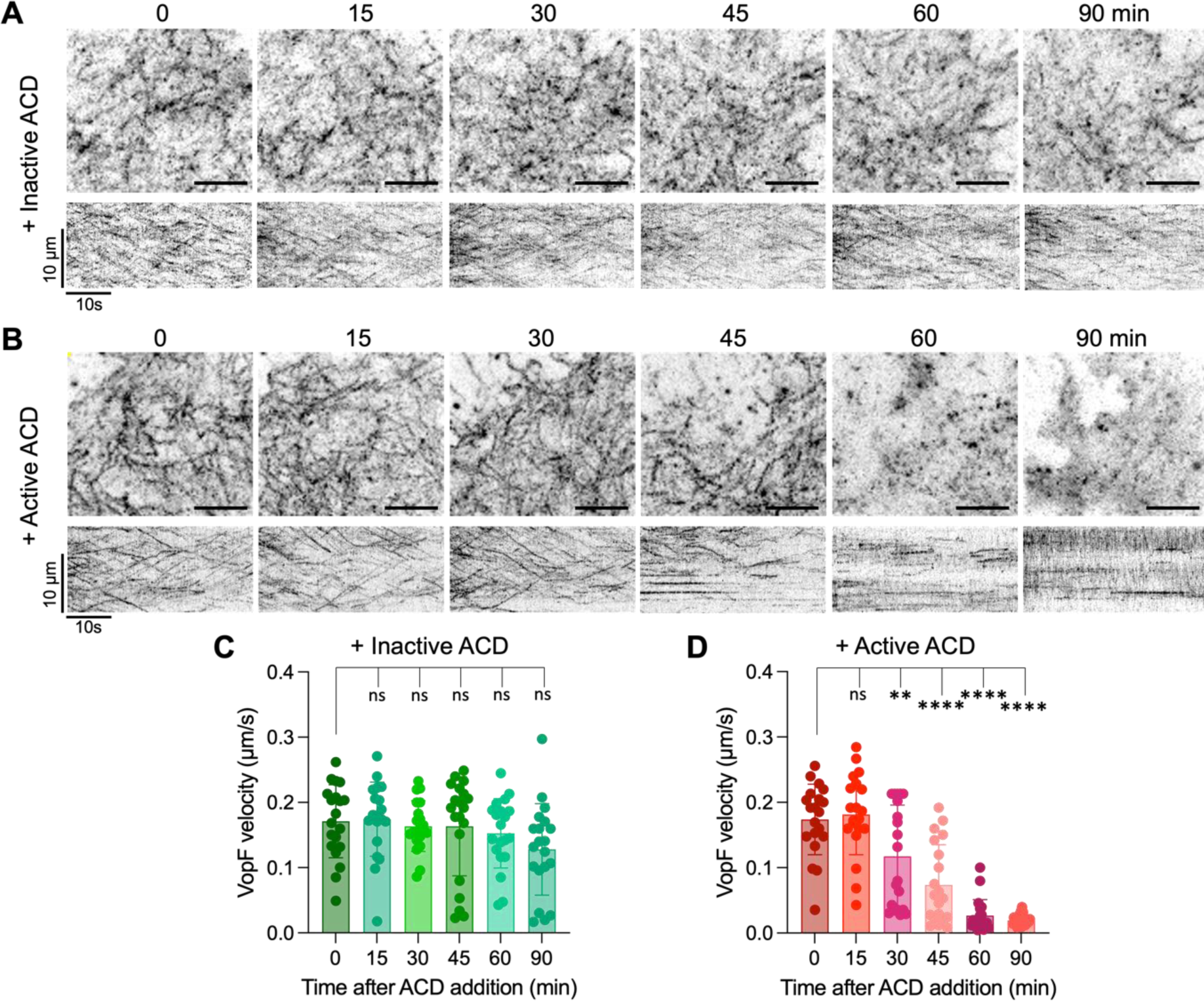
ACD inhibits pointed-end actin elongation activity of VopF in living cells. (A,B) Pointed-end actin elongation activity of VopF was visualized by SiMS microscopy of XTC cells expressing low single-molecule level of EGFP-VopF. Maximum intensity projections (MIP) and kymographs (kymo) of time-lapse images are shown after treating cells with a mixture of PA (2.5 nM) and either active (A) or inactive (B) LF_N_-ACD (1 nM); time is shown in minutes. Scale bars for all MIP images are 5 µm. (C,D) VopF speckle velocities were measured using kymograph analysis and plotted as means (n=20) ± SE at the indicated time point after the ACD addition. Ordinary one-way ANOVA with Dunnett’s multiple comparisons test was performed to determine statistical significance; ns, not statistical; ** = 0.0057; **** < 0.0001.

### Expression and secretion of RtxA and VopF toxins by *V. cholerae* strains

In an attempt to understand whether the expression of the two competing toxins, VopF and ACD-containing MARTX, is reciprocally coordinated to avoid potentially counter-productive interference, we assessed the expression of both toxins in the *V. cholerae* AM-19226 strain. To mimic an environmental cue used by *Vibrio* and other bacteria to recognize their location in the digestive tract and promote virulence gene expression^44^, *V. cholerae* strains bearing *lacZ* transcriptional reporter fusions to MARTX/T1SS and VopF/T3SS encoding genes were grown in the absence and presence of bile. As positive controls for bile induction, reporter fusions to operons encoding the T3SS structural genes *vcsRTCNS2* and *vcsVUQ2* were included^45^. We observed over 7-fold higher expression of both operons in the presence of bile, compared to when cells were grown in LB alone, consistent with previous reports^45,46^. We found that bile also induced the expression of the T1SS structural gene *rtxB* and the ACD-encoding *rtxA* gene by more than 4-fold (Fig. 4). However, the expression of *vopF* was not only notably weaker, but also did not substantially change in response to bile (Fig. 4). These data suggest that expression of the genes encoding the the RTX toxin and VopF proteins is likely regulated independently by different environmental signals. Thus, transcriptional control of effector expression may be one level by which the pathogen avoids interference when encoding both toxins, although other mechanisms are possible and remain to be established.

**Figure 4.**
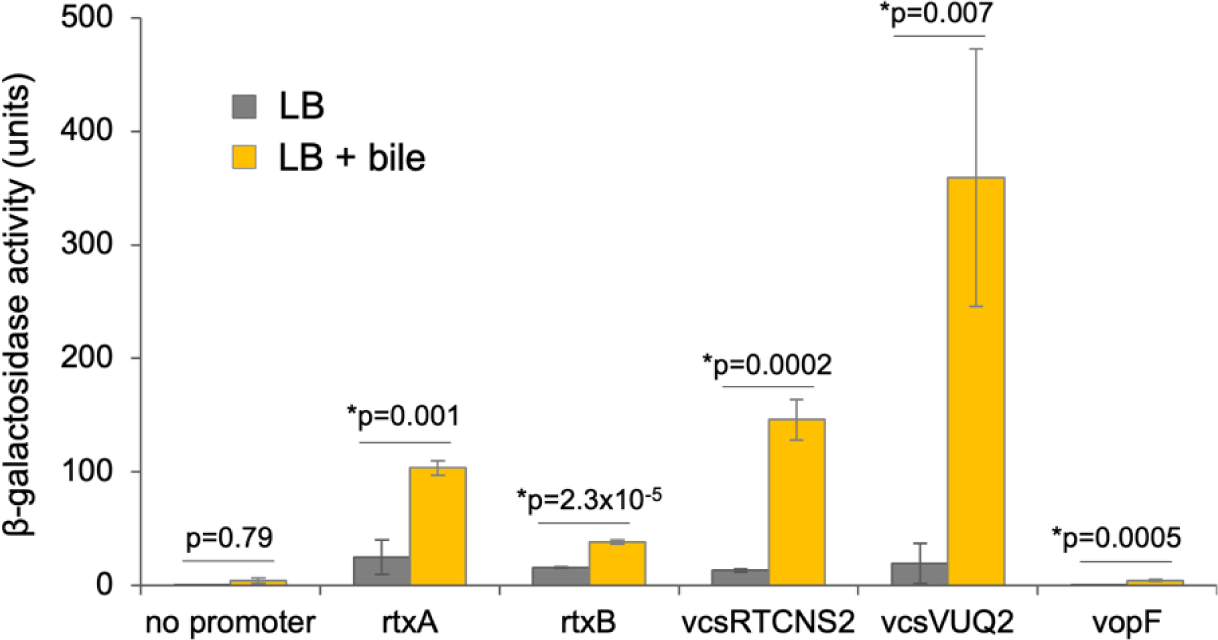
Expression of *V. cholerae* T1SS and T3SS structural genes, ACD-encoding rtxA, and VopF genes in the presence and absence of bile. *lacZ* transcriptional reporter fusions carried in single copy in the AM-19226 strain background are indicated on the *x*-axis: *rtxA* and *rtxB* indicate fusions to the promoter region of T1SS related genes, whereas T3SS genes include the structural gene operons *vcsRTCNS2* and *vcsVUQ2* and vopF, which encodes a T3SS translocated effector. Three colonies from each strain were grown overnight at 37°C in LB media supplemented with 0.4% bile, and beta-galactosidase activity was measured using a standard kinetic assay, where units equal micromoles of ONPG hydrolyzed per minute per OD_600_. A promoterless-lacZ fusion strains served as the negative control and exhibited less than 20 units activity. The experiment was repeated with similar results. Data is presented as means (n=3) ± SD. ANOVA single factor followed by Student’s t-test with Bonferroni correction for multiple comparisons was performed to determine statistical significance.

## Discussion

While achievable in experimental animals^47^, co-infection of the human intestinal tract with two or more pathogenic bacteria is infrequent^5,48^. In contrast, virus-virus, virus-protozoan, and virus-bacterial infections are much more common^5^, likely because the competition between distinct pathogens is much less intense than that between similar ones with substantially overlapping niches and survival strategies. The analysis of the intense competition between bacterial species is often reduced to the most evident scenario of competition for resources (exploitative competition) and direct eradication of the competitors by toxic molecules (interference competition)^6^. While the role of the host’s ability to regulate microbial communities via immune mechanisms is well recognized, the ability of bacteria to manipulate the host to gain a competitive advantage is less obvious, difficult to study in complex *in vivo* systems and, likely for these reasons, not commonly discussed in the literature. Pathogenic bacteria employ a range of bacterial effector proteins and toxins to invade host cells or establish a robust intestinal colonization while evading immune responses. The combinatory actions of these proteins are determined by their targeting specificity, the temporal control of their production and release, and the potential synergy or interference in their activities. Interestingly, effector proteins produced by different pathogens, or even different strains of the same pathogen, may exert antagonistic effects on host cell machinery.

In this work, we utilized *Salmonella* invasion into cultured epithelial and macrophage cells as a model for testing the hypothesis that the competition between pathogenic bacteria can be mediated via their antagonistic effects on host systems essential for pathogenesis, *e.g.,* the actin cytoskeleton. To reduce the complexity, we simulated the actin-targeting effects of various bacteria by using only their effector proteins and delivering them via a modified anthrax toxin machinery. We intentionally limited our focus strictly to the actin cytoskeleton by utilizing actin-specific toxins and effector proteins whose mechanisms are reasonably well understood. Thereby, we evaluated the potential competition between different actin-specific effector proteins from the same or different pathogens that disturb the actin cytoskeleton via distinct mechanisms.

As some effector proteins hijack the actin motor force for invasion or locomotion inside the host cell and others disrupt the actin integrity to compromise the protective function of immune and epithelial cells, we anticipated that these two groups of effectors are likely to interfere with each other. Indeed, we observed various degrees of inhibition of *Salmonella* invasion into macrophage and epithelial cells pre-treated with the effectors causing actin depolymerization (SpvB), compromising the integration between cytoskeletal and cellular elements (TccC3), inactivating Rho GTPases (RID), or competitively inhibiting F-actin assembly factors (actin oligomers produced by ACD) (Fig. 1). These effector proteins disrupt the actin cytoskeletal structure and dynamics in various, independent ways, thus they would be predicted to interfere with invasion orchestrated by *Salmonella* SPI-1 effector proteins.

Invasion of *S.* Typhimurium into polarized enterocytes relies on the production of SipA^49^, the effector that stabilizes filamentous actin^50–53^ and promotes membrane fusion events^54^. Invasion into non-polarized cells requires the combined action of SopB, SopE, and SopE2 effectors^55–58^, which coordinate the activation of small GTPases to promote actin-dependent membrane ruffling driving the engulfment of the pathogen into non-polarized cells. While non-compatible effects of effector proteins from different pathogens are not surprising (either coincidental or serving competition purposes), antagonistic effects from effectors of the same pathogen are also not rare. Thus, *Salmonella* SipA and SpvB effectors are engaged in mutually counter-productive stabilization and destabilization of actin filaments, respectively. A similar conflict can be expected between SpvB and SopB/SopE effectors acting upon Rho GTPases^59^.

Given that the bacteria’s own effector proteins can inhibit or even cancel each other’s activities, it is imperative that their effects should be separated either spatially (*e.g.,* by acting upon different cell types or different subcellular compartments) or temporally. The latter case applies for separating the actin-dependent invasive activity mediated by the SPI-1 effectors (*e.g.,* SipA, SopB, SopE) from the actin-disrupting activity of SpvB secreted via SPI-2 after *Salmonella* has invaded a host cell and established within the replicative, perinuclear niche in the *Salmonella*-containing vacuole^60^. When properly separated, the antagonistic effects may even be beneficial. Thus, we speculate that ADP-ribosylation of actin by SpvB may be important to prevent invasion of other *Salmonella* into the same cell, limiting the intraspecies competition for host cells. Notably, this inhibition is more relevant to *Salmonella* invasion of polarized enterocytes, which occurs through a single bacterium “discreet” entry^49^, as opposed to the cooperative engulfment of multiple pathogens into non-polarized epithelial cells^61^, not found in the intestine. The application of this mechanism of “discrete” invasion allows for the avoidance of intracellular competition between the same species of bacteria. Whether SpvB can reduce reinfection of already infected cells *in vivo* remains to be established. If confirmed, this mechanism may contribute to the recognized role of this effector in the systemic spread of non-typhoidal *Salmonella* strains and colonization of vital organs^62^.

While SpvB is a *Salmonella* effector, it is a member of a large group of actin-specific mono-ADP-ribosylating toxins (mARTs), broadly expressed in the microbial world^8,63–65^. In the context of the proposed interspecies competition mediated via host systems, release of actin-targeting mARTs can interfere with the bacteria that benefit from robust actin dynamics in the host cell.

Another pair of effector proteins with conflicting effects on actin are *V. cholerae* ACD and VopF. Actin reorganization by the VopF/VopL effectors produced by *V. cholerae* and *V. parahaemolyticus*, respectively, are required for intestinal colonization using animal models of infection^37–39^. While the detailed mechanisms of colonization remain to be understood, at the cellular level, VopF compromises the integrity of epithelial tight junctions^38^ and VopL perturbs the membrane localization of innate immune enzymes (*e.g.,* NADPH oxidase^39^). At the molecular level, these effects result from robust nucleation and unconventional pointed-end processive elongation of actin filaments (the latter of which is prohibited under the physiological cellular conditions), leading to disruption of the cellular actin network polarity^41^. With their three tandem-organized G-actin binding WH2 domains, which are required for the above activities, VopF/L are canonical targets for the ACD-produced actin oligomers^15^, as confirmed by the oligomers’ ability to potently inhibit nucleating, severing, and polymerizing activities of VopF/L (Figs. 2 and 3).

As part of a giant multifunctional MARTX toxin, ACD is secreted to the environment via T1SS, enabling long-range targeting of any cell, immune or epithelial, that has a respective receptor, as demonstrated in cell culture experiments^66,67^. While the individual effector domains included in MARTX toxins varies in different strains of *V. cholerae* and related bacteria, VopF is solely a T3SS effector protein, delivered by an independent mechanism^21,25^. If both ACD and VopF were MARTX effector domains in the same molecule, their conflicting activities would be co-localized in both space and time, and thus antagonistic by default. Instead, T3SS effector VopF is injected upon contact with the host cell and, therefore, is more likely to directly benefit the secreting bacterium, rather than causing global effects. Whether this and other similar conflicts are reconciled by controlling the expression of such effectors at different stages of infection or due to their cellular and subcellular specificity remains unknown. In the case examined here, the two genes appear to be regulated independently: while bile enhanced *rtxA* expression, *vopF* expression remained low in the presence of bile and notably lower overall, suggesting a different environmental cue promoting *vopF* expression (Fig. 4). It is tempting to speculate that *vopF* expression may respond to direct contact with the host cells, or a temporal signal in the intestine following bile exposure.

The proposed possibility for indirect bacterial competition is not limited in our opinion to antagonistic effects of bacterial effector proteins on the actin cytoskeleton. Competitive pressure of commensal bacteria is a major mechanism protecting the digestive tract from pathogenic species. While diarrhea can be initiated by the host and have a protective function, at least some of the diarrhea-causing bacteria (*e.g.*, *V. cholerae* and *Salmonella* strains) benefit from this induced imbalance in the host’s water-electrolyte homeostasis not only by release of nutrients and a facilitated evacuation and spread, but also by drastically reducing the complexity of the microbiome^68^, reducing the competitive pressure from commensal microorganisms.

To summarize, in this study we explored the concept that competition in bacterial communities can be mediated via a functional interference of their effects on the host. Our focus solely on an *in vitro Salmonella* invasion model as a function of the host’s actin cytoskeleton perturbed by different effector proteins allowed us to confirm that the proposed competition mechanisms are relevant at least for simplified systems, opening the possibility of their broader applicability to host-associated bacterial communities. Tentatively, when applied to organisms within the same species, this mechanism may boost the efficiency of infection by preventing the invasion of the same cell by multiple bacteria. The proposed mechanism may also account for the low likelihood of effective co-infection of the digestive tract with two or more bacterial pathogens simultaneously.

## Materials and Methods

### Cell culture infection

J774A.1 and HeLa cells (ATCC) were grown until confluent in surface-treated tissue culture flasks (Fisherbrand) in Dulbecco’s modified Eagle medium (DMEM, Gibco-Life Technologies) supplemented with 10% heat-inactivated fetal bovine serum (FBS, Corning) and penicillin-streptomycin (Gibco-Life Technologies) at 37°C in a humidified incubator with 5% CO_2_. To remove any dead cells, the tissue culture flasks were washed with PBS, prior to harvesting the cell monolayer with 0.25% trypsin-EDTA (1X) (Gibco-Life Technologies). Cells were then washed to remove trypsin, counted using trypan blue (Gibco-Life Technologies) exclusion, and seeded at a density of approximately 1.0 × 10^5^ cells/well in 24-well polystyrene microplates (Falcon) for infection studies.

For invasion studies, overnight cultures of wild-type *S*. Typhimurium strain 14028 were grown in LB medium. The next day, the culture densities were adjusted to an OD_600_ of 0.4 using DMEM. Prior to addition of the bacteria, J774A.1 and HeLa cells were incubated in DMEM supplemented with 10% FBS in the absence or presence of a mixture of 2.5 nM PA with 1 nM LF_N_-toxins for 1 hr at 37°C in a humidified incubator with 5% CO_2_. Then, bacteria were added to the toxin-pretreated cells at a multiplicity of infection (MOI) of 10 in 1 mL DMEM supplemented with 10% FBS. The plates were centrifuged at 1000 rpm for 10 minutes to synchronize the infection. After 1-h incubation, extracellular bacteria were removed by adding 100 mg/mL gentamicin (Gibco-Life Technologies) for 1 hr followed by washing with phosphate-buffered saline (PBS). The infected cells were lysed with 1% triton X-100 (Calbiochem) for 15 min. The cell lysates were then serially diluted, plated onto LB agar, incubated at 37°C overnight and enumerated for CFU/mL and percent invasion (bacteria recovered / bacteria infected).

### Protein purification

Actin was purified from rabbit skeletal muscle acetone powder (Pel-Freez Biologicals) as reported^69^. *Bacillus anthracis* protective antigen (PA) was expressed in *E. coli* purified from the periplasmic space^70^. Actin-targeting constructs of the effector domains ACD and RID from *V. cholerae* MARTX toxin and TccC3-hvr (hypervariable region) effector domain of Tc toxin from *P. luminescens* C-terminally fused to the N-terminal domain of *B. anthracis* lethal factor (LF_N_-ACD, LF_N_-RID, and LF_N_-TccC3) in-frame with an N-terminal 6-His tag were expressed and purified as previously reported^23,27,71^. An LF_N_-fusion construct of the catalytic domain of *S.* Typhimurium SpvB (residues 367-591) was similarly cloned into a modified^72^ pCold(I) plasmid (Takara). The VopF fragment encoding amino acid residues 129-530 containing the three WH2 domains and the VopF C-terminal actin-binding domain (VCD) was synthesized by GenScript® (Piscataway, NJ). VopL (amino acids 90-484) cDNA in pTYB12 was a gift of Roberto Dominguez (Perelman School of Medicine-University of Pennsylvania)^73^. VopF and VopL cDNAs were cloned into the pCold(I) vector for recombinant protein purification from *E. coli* and into dCMV-EGFP for SiMS microscopy experiments as described^41^. The recombinant proteins were expressed in *E. coli* and purified using either Talon cobalt resin (Clontech) or HisPur cobalt resin (ThermoFisher). ACD-crosslinked actin oligomers were prepared using thermolabile ACD from *Aeromonas hydrophila* (ACD*_Ah_*)^74^ as desribed^16^. Recombinant human PFN1 was purified as described^75^.

### Bulk pyrene-actin polymerization assay

Bulk pyrene-actin polymerization assays were set up as previously described^15,16^. Inhibitory activity of the oligomers on actin nucleation was measured by following the protocol described previously^15,76^. Briefly, the points from 5-20% maximal fluorescence of each pyrene fluorescence trace were fit to a quadratic equation:

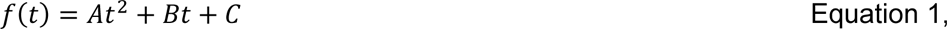

where *A* is dependent upon the nucleation rate, *B* is dependent upon the number of barbed ends and *C* is dependent upon the concentration of actin. By using initial points where few filaments have formed in the presence of profilin, the values for *B* and *C* were considered equal to zero.

The nucleation rate (*NR*) is then determined using the following equation:

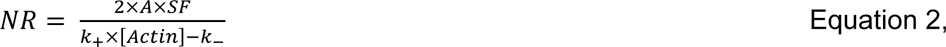

where *k_+_*is the rate of barbed end elongation (11.6 μM^−1^s^−1^) and *k_-_* is the rate of pointed end depolymerization (1.4 s^−1^) at 22 °C^77^, *A* is determined from the equation (1), *[Actin]* is the initial concentration of actin monomers, and *SF* is the actin filament concentration scaling factor determined by subtracting the critical concentration of actin polymerization (0.1 μM^77^) from *[Actin]* and dividing by the total fluorescence change.

The normalized inhibitions (F/ΔF) of calculated nucleation rates were fit to a binding isotherm:

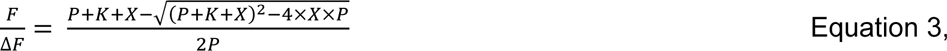

where *P* is the concentration of functional units of VopF/L, and *X* is the concentration of actin oligomers, and *K* is the apparent *k_d_*value.

### Total internal reflection fluorescence (TIRF) microscopy

*In vitro* TIRF microscopy experiments were performed as described^15,16^. For inhibition of nucleation activity, rabbit skeletal actin (20% Oregon Green-labeled or Alexa 488-labeled; 1.0 μM final concentration) was switched from Ca^2+^-ATP to Mg^2+^-ATP state by incubation with exchange buffer (final composition: 50 μM MgCl_2_ and 0.2 mM EGTA) for 2 min. Actin was added to the mixture of 10 nM VopL and PFN1 in the following final buffer composition: 10 mM imidazole, pH 7, 50 mM KCl, 50 mM DTT, 1 mM MgCl_2_, 1 mM EGTA, 0.2 mM ATP, 50 μM CaCl_2_, 15 mM glucose, 20 μg/mL catalase, 100 μg/mL glucose oxidase, 3% glycerol, and 0.5% methylcellulose-400cP (Sigma Aldrich). Immediately after adding actin, samples were transferred to a NEM-myosin treated flow chamber^78^ and images were collected for 8 min, at 5-s intervals. Images were collected on a Nikon Eclipse Ti-E microscope equipped with a perfect focus system, through-the-objective TIRF illumination system, and DS-QiMc camera (Nikon). The filaments formed were manually counted at 8 min using Fiji/ImageJ software^79^.

The number of filaments formed (*N(C)*) in the presence of each concentration of the oligomers were fit to an exponential decay:

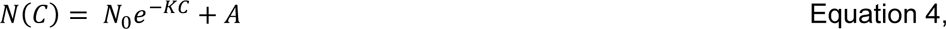

where *N_0_*is the number of filaments nucleated above spontaneous nucleation in the presence of VopL, *C* is the concentration of the oligomers, A is the number of filaments formed based on the nucleation rate of actin alone, and *K* is the IC_50_ value.

For inhibition of filament severing, Mg^2+^-ATP actin was prepared as above. After most filaments had grown to more than 10 μm in length, 10 nM VopL and varying concentrations of actin oligomers were flowed into the chamber to replace the solution of free actin monomers. To determine the efficiency of severing by proteins in the presence of the oligomers, filaments lengths were measured in the frame prior to the addition of the oligomers and the protein of interest and manually counting the number of severing events for 60 frames or 300 s using Fiji software.

### Single-molecule speckle (SiMS) microscopy

*Xenopus laevis* XTC cells were grown in 70% Leibovitz’s L-15 medium at 23°C without CO_2_ equilibration as described previously^80^. The cells were transiently transfected with the dCMV-EGFP-VopF construct were seeded onto poly-D-lysine-coated coverslips in Attofluor chambers (ThermoFisher Scientific) and imaged using TIRF module on Nikon Eclipse Ti-E inverted microscope equipped with perfect focus system, Nikon CFI Plan Apochromat λ x100 oil objective (NA 1.45), and iXon Ultra 897 EMCCD camera (Andor Technology) as described^41^. Cells were treated with mixtures of either active or inactive (EE1990,1992AA)^15,43^ LF_N_-ACD complexed with PA (at 1 and 2.5 nM final concentrations, respectively) and the intracellular motility of EGFP-VopF speckles was recorded using time-lapse imaging every 0.5 s for 2 min with 13 min intervals. VopF velocities were quantified using kymograph analysis as described^41^.

### Construction of the *rtxA::lacZ* and *rtxB::lacZ* reporter strains and β-galactosidase assay

The *rtxA::lacZ* and *rtxB::lacZ* reporter fusions were constructed in pEH3 vector using standard methods as previously described, and integrated in single copy into the *V. cholerae* strain AM-19226 *lacZ* locus^45^. To determine the relative expression level of each construct, three colonies from each strain were grown in either LB broth or LB broth with 0.4% bile overnight at 37°C and β-galactosidase activity assayed as previously described^45^. Briefly, the cultures were subjected to centrifugation, and the resulting pellets were re-suspended in Z-buffer^45^ with β-mercaptoethanol. The OD_600_ was measured, and the cell suspension was used to determine the cleavage of ortho-nitrophenyl-β-galactoside (ONPG) as measured by the OD_420_ in a standard kinetic β-galactosidase assay^81^. The β-galactosidase activity was calculated as the µM of ONP formed per min per cell using the formula:

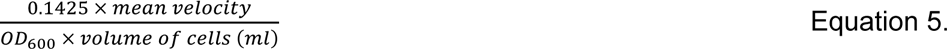

## Movie Legends

**Movie 1. Directional movement of VopF due to its pointed-end actin elongation activity is not impaired by the inactive ACD.** A multi-stack montage of time-lapse images of a peripheral region of an individual XTC cell expressing low levels of EGFP-VopF and treated with PA and inactive LF_N_-ACD EE1990,1992AA mutant (at 2.5 and 1 nM final concentrations, respectively). Images were collected using TIRF microscopy at 0.5 s intervals for 2 min for each 15-min time point (only the first 1.5 min is shown for each) with 13 min intervals between the adjacent time points. Scale bars are 5 µm.

**Movie 2. Active ACD inhibits VopF pointed-end actin elongation activity.** A multi-stack montage of time-lapse images of a peripheral region of an individual XTC cell expressing low levels of EGFP-VopF and treated with PA and active LF_N_-ACD (at 2.5 and 1 nM final concentrations, respectively). Images were collected using TIRF microscopy at 0.5 s intervals for 2 min for each 15-min time point (only the first 1.5 min is shown for each) with 13 min intervals between the adjacent time points. Scale bars are 5 µm.

## Acknowledgements

We thank Jenna Hindsley for her contribution to the beta-galactosidase experiment. The work was supported by NIH R01 GM114666 and NIH R01 AI126005 grants (to DSK and MD, respectively), by OSU PHPID fellowship (to DBH), and funds provided by the Infectious Disease Institute (idi.osu.edu) and Nationwide Children’s Hospital (to JG).

